# Re-establishment of TAD boundary organization during DNA replication

**DOI:** 10.64898/2026.07.08.737321

**Authors:** Theodore Busby, Adib Keikhosravi, Mohamadreza Fazel, Faisal Almansour, Kathleen S. Metz Reed, Laurent Ozbun, Tatiana Karpova, Gianluca Pegoraro, Tom Misteli

## Abstract

A ubiquitous feature of higher order genome organization is the presence of topologically associating domains (TADs). The chromatin architectural proteins cohesin and CTCF are known critical organizers of TADs, but the mechanisms of TAD establishment and maintenance, including their accurate duplication during genome replication, are not well characterized. To address this gap, we used high-throughput imaging-based CRISPR/Cas9 knock-out screening to discover chromatin factors involved in maintenance and establishment of TADs. Among the cellular factors that affect TAD organization, we found enrichment for cell cycle proteins, especially components of the DNA replication machinery. Accordingly, we demonstrate that TADs undergo temporary unfolding during S-phase DNA replication and that interference with progression through replication impedes restoration of normal TAD structure. Mechanistically, inhibition of the RPA complex prevents cohesin and CTCF binding, delays post-replication TAD re-folding and affects TAD folding in non-cycling cells. These results provide novel insights into how TAD structures are re-established during genome duplication.

## Introduction

Chromatin is spatially organized in three dimensions in the cell nucleus^1^. A major organizational feature of the genome is folding of the chromatin fiber into topologically associating domains (TADs). These domains are typically several hundred kilobases to mega-bases in size and are defined as genome regions which preferentially interact with each other rather than their neighboring sequences^1,2^. TADs are thought to be functionally relevant by promoting high frequency interactions, such as between promoters and enhancers within a TAD, while disfavoring cross-contacts with neighboring domains^3–5^. TADs are delineated by flanking boundary elements that contain binding sites for the chromatin protein CTCF (CCCTC-binding factor) and for the ring-shaped cohesin ATPase complex, consisting of SMC1A, SMC3 and RAD21^1,6,7^. The organization and integrity of TADs are associated with regulation of developmental gene expression^8–11^, genome stability^12^, replication and the cell cycle^13–17^. Despite their mappability by population-based biochemical methods, such as Hi-C, single-cell analysis, and live-cell imaging have demonstrated that TADs are dynamic in nature with their boundaries undergoing rapid cycles of pairing and unpairing on the scale of minutes^18,19^.

The prevailing model for TAD formation is through a mechanism referred to as loop extrusion^1,2,20–22^. In this process, the ring-like cohesin complex associates with chromatin via the loading factors NIPBL and MAU2^23,24^. The ATPase motor activity of cohesin then pulls chromatin through the ring until cohesin encounters bound CTCF at TAD boundaries, thereby forming a loop^1,2,7^. The cohesin complex is then removed from chromatin by the release factor WAPL^7,24^. The constitutive activities of the cohesin complex can be modified in specific cellular contexts. For example, recruitment of cohesin to tissue-specific genomic sequences is modulated by the cohesin subunit STAG2^25,26^. Furthermore, acetylation of SMC3 by the ESCO acetyltransferase blocks the release of cohesin by WAPL, stabilizing the complex on chromatin^27–29^. In contrast, the GSK3 kinase recruits WAPL to release cohesin from chromatin^30^. Dysregulation of these mechanisms has physiological consequences leading to pathologies like cancer and cohesinopathies^31^. Similarly, regulation of CTCF via recruitment and post-translational modifications also contributes to organization of the chromatin fiber^8,32–35^. The growing list of regulatory factors that affect chromatin organization highlights the complexity of the mechanisms involved in establishing and maintaining TADs, including during the cell cycle.

In this study, we sought to systematically identify novel cellular factors that determine TAD architecture. To do so, we measured the physical distance between boundaries of individual TADs at the single-allele level by high-throughput DNA FISH (Fluorescence In Situ Hybridization) using the *MYC* locus on chromosome 8 as a model system. Amongst other factors, we identify the replication protein A (RPA) complex as a prominent cell cycle regulator of TAD structure. Tracking of TAD boundaries through S-phase demonstrated their physical separation concomitant with their replication. Consistent with the role of RPA in TAD duplication, inhibition of RPA led to separation of TAD boundaries in a cell-cycle dependent manner, and blocked replication-associated boundary separation and prevented loading of cohesin. Taken together, our results identify novel nuclear factors involved in determining TAD organization and provide new insight into how higher order chromatin structures are re-established during replication.

## Results

### A high-throughput imaging pipeline to visualize TAD boundary architecture

We sought to identify novel nuclear factors involved in establishment and maintenance of TAD architecture using an unbiased CRISPR-based high-throughput imaging screen. As a model, we selected the *MYC* TAD on chromosome 8 in the HCT116 colorectal cancer cell line (Fig. 1A). The *MYC* TAD is ideal for this approach because of its relatively large size of approximately 3 Mb, extensive sub-TAD structure, and the *MYC* proto-oncogene being the only protein coding gene expressed from this TAD^5,36–38^. To assess TAD boundary architecture, we measured the distances between the 5’ and 3’ TAD boundaries at single alleles using 2-color high-throughput fluorescence in situ hybridization (HiFISH)^18,39–41^ (Fig. 1B). To achieve highest accuracy in TAD boundary distance measurements, we determined the center-to-center distances of FISH signals of typically 2,000-8000 individual alleles on projected 3D image stacks using a previously characterized custom-built image analysis platform^42^ and a euclidean distance measurement algorithm^41^ (Fig. 1C, D). As expected, based on the known dynamic behavior of TAD boundaries^19^, *MYC* TAD boundary distances varied amongst individual alleles in the population typically from 150 nm to ∼1 µm with a mean distance of ∼400nm in control cells (Figure 1C-D). As a proof-of-principle for our ability to detect changes in boundary distances, we performed acute depletion of the cohesin subunit RAD21 in HCT116-RAD21mAID degron cells, which has previously been shown to cause loss of TAD organization^36^ (Extended Data Fig. 1A-B). Depletion of RAD21 for up to 6 hours indeed increased the distance between the boundaries from a mean of ∼500 nm to ∼950 nm (p-value: 1x10^-6^; Extended Data Fig. 1B). Consistent with a role for RAD21 in boundary pairing during loop extrusion, RAD21 loss had a more pronounced effect on the *MYC* TAD boundaries as compared to a control region containing an equidistant non-boundary genomic region (Extended Data Fig. 1C-D). Taken together these results establish a robust assay to assess TAD boundary distances for use in high-throughput imaging-based screening.

**Fig. 1.**
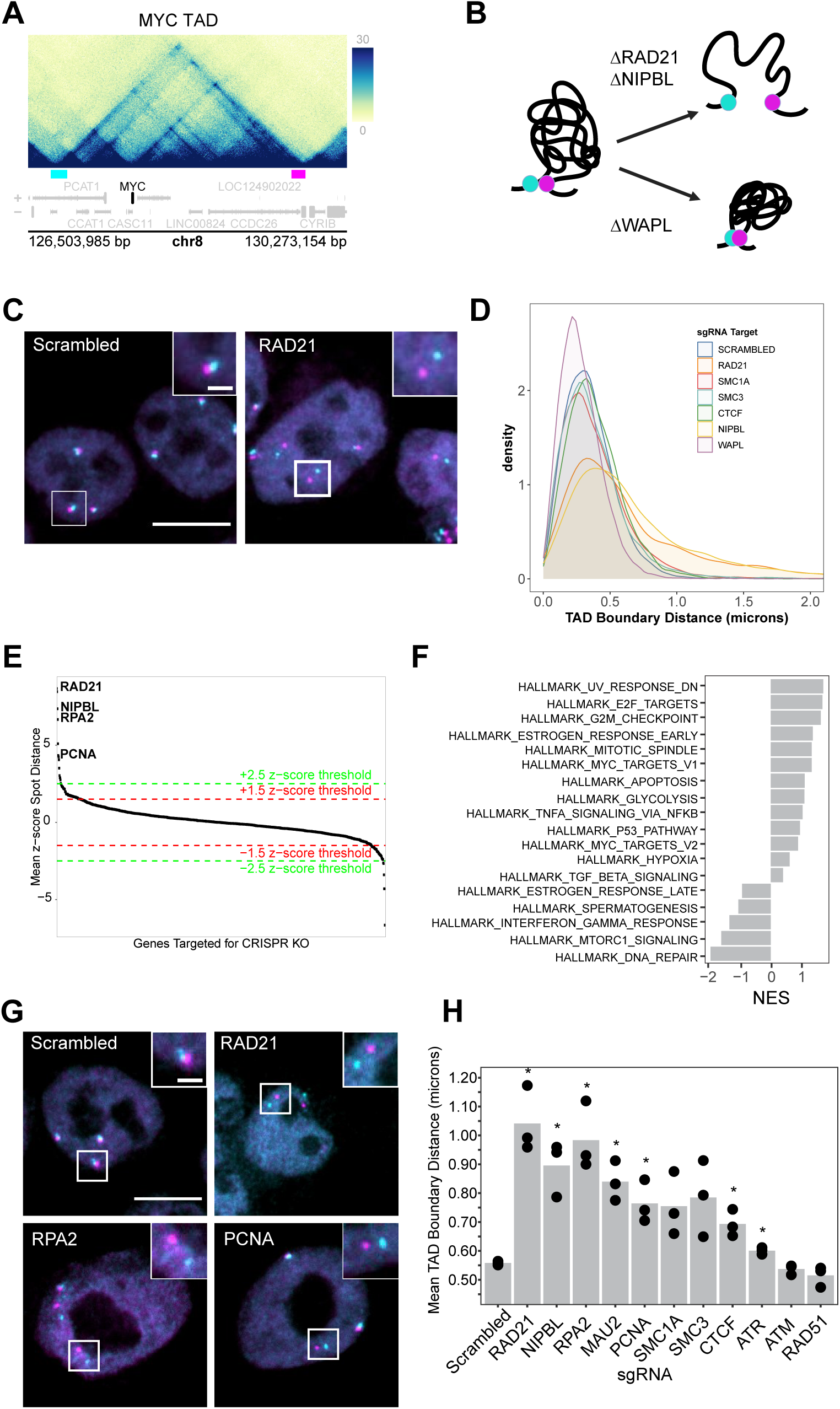
CRISPR-KO screen identifies regulators of TAD boundary proximity. **A**) HiC map of the *MYC* TAD with location of DNA FISH probes (cyan = 5’ probe, magenta = 3’ probe) in HCT116 cells (4DN Data Portal: 4DNES3QAGOZZ). **B)** Model depicting TAD structure and consequences of cohesin deletion. FISH probes are indicated in cyan and magenta. **C)** Representative images of MYC TAD boundary FISH after RAD21 knockout by CRISPR. 5’ TAD boundary (cyan), 3’ TAD (magenta); scale bar: 10µm. **D**) HiTIPs analysis of boundary distances after cohesin and CTCF knockout. Density curves represent the distribution of TAD boundary distances across the population of alleles measured. Typically 2,000-8,000 alleles were measured per experiment. **E)** Rank of z-scores for *MYC* TAD average boundary distances of all 1064 screened CRISPR target genes based on three biological replicates with +/-2.5 and +/-1.5 z-score cut-offs. See Extended Data Table 1 for numerical values. **F)** Enrichment of cellular processes for hits with z-scores over 1.5. **G)** Representative images of replication-related hits identified in the screen. scale bar: 10µm. **H)** Average TAD boundary distances (bar) for three biological replicates (dots) for top hits. Statistical analysis performed by two-tailed t-tests; *P< 0.05.

To establish a pipeline for CRISPR knock-out (CRISPR-KO) high-throughput screening, we performed a pilot screen to target several cohesin subunits and CTCF in HCT116-Cas9 cells. Oligo sgRNAs were introduced by reverse transfection for 72 hours in 384-well imaging plates for CRISPR-KO gene disruption, and their effect on the *MYC* TAD boundary distances was measured by automated HiFISH (Fig. 1C, D). Depletion was efficient as sgRNA targeting the *LMNA* gene led to robust knockout of lamin A and C protein in ∼75% of cells, as determined by immunofluorescence imaging (Extended Data Fig. 1E). Consistent with the acute depletion via degron, CRISPR-KO of RAD21 led to separation of *MYC* TAD boundaries, increasing the mean boundary distance from ∼400 nm to ∼750 nm (Fig 1C-D, p-value: 1x10^-4^). Similarly, CRISPR-KO of the cohesin loader NIPBL also led to separation of the TAD boundaries to ∼750 nm with a large shift in the distribution towards longer distances at the allelic level (Fig. 1D, Extended Data Fig. 1F; p-value: 1x10^-4^). We also observed a shift in the boundary distance distribution for CRISPR-KO of cohesin ring subunits SMC1A (p value: 1x10^-4^) and SMC3 (p value: 1x10^-4^) as well as for the architectural protein CTCF (p value: 0.016) (Fig. 1D). CRISPR-KO of the cohesin unloader WAPL, on the other hand, led to shorter center-to-center mean distances from ∼400 nm to ∼350 nm (p value: 1x10^-4^) between the *MYC* TAD boundaries as expected based on WAPL’s role in releasing cohesin from looped boundaries (Extended Data Fig. 1F)^24,43^. These results demonstrate the feasibility of using sgRNA-mediated knockdown as an approach to identify cellular determinants of TAD organization and they identify CRISPR-knockout of RAD21 and NIPBL as strong positive controls for a high-throughput screen.

### Identification of novel modulators of TAD architecture

We applied this high-throughput imaging pipeline to screen a library of sgRNAs targeting 1,064 genes that express nuclear localized proteins, especially chromatin proteins, epigenetic modifiers, transcription factors, and cell cycle regulators (See Extended Data Table 1 for full library). HCT116-Cas9 cells were reverse transfected in a 384-well format for 72 hours with three sgRNAs per gene in individual wells (see Methods). *MYC* TAD boundary distances were measured per target gene to generate population means per sgRNA target and normalized across all distances measured per assay plate, including control wells and targeted sample wells, to generate a statistical z-score based on the average distances measured for all sgRNA targets (see Methods). We performed the screen in triplicate with scrambled sgRNA as a negative control, sgRNA against the essential *PLK1* gene as a transfection efficiency control indicated by cell death, and sgRNAs against *NIPBL* and *RAD21* as positive controls (see Extended Data Table 1). We identified 11 genes (∼1% of library) that resulted in significant changes to TAD boundary distance with a z-score cutoff of > 2.5 and 39 genes (3.5% of library) with a z-score cutoff > 1.5. Reassuringly, of the top hits, five genes were known architectural TAD factors including the cohesin subunits RAD21, SMC1A, and SMC3 (Fig. 1E). Cohesin loaders NIPBL and MAU2 were also identified as significant hits (Fig. 1E), confirming the validity of our screening approach. A set of 48 targets were orthogonally validated by siRNA knockdown (Extended Data Table 2).

Amongst the top screen hits, we observed an enrichment of general chromatin organizers (SMCs, TOP2A), cell cycle regulators (CDC45, SKA2) and in particular proteins involved in replication including PARP3, ORC2 and PCNA (Fig. 1F, Extended Data Table 1). In fact, Replication Protein A 2 (RPA2) and Proliferating Cell Nuclear Antigen (PCNA) were amongst the highest ranked hits (Fig. 1E, G; Extended Data Table 1). Compared to scrambled controls, knockout of each of these genes caused a significant increase in TAD boundary distance from ∼500 nm in the control to ∼980nm (p value = 0.025) and ∼760 nm (p-value = 0.040) in sgRPA2 and sgPCNA targeted cells, respectively (Fig. 1G-H, Extended Data Table 1), comparable to knockout of NIPBL (Fig. 1G-H). These effects were not due to the known role of RPA2 in the DNA damage response (DDR) because several DDR factors represented in the library, including ATM, ATR and RAD51, were not hits or had an opposite effect on TAD boundary distances than RPA2 (Fig. 1E, H, Extended Data Table 1, Extended Data Fig. 2B)^44^. The strong preponderance of replication-related factors prompted us to examine the role of replication and of RPA2 in regulating TAD boundary architecture in more depth.

### The RPA complex regulates TAD boundaries

RPA2 is the 32kDa subunit of the heterotrimeric replication protein A (RPA) complex that binds single stranded DNA (ssDNA) at the replication fork during DNA synthesis^45^ (Fig. 2A). Because the other RPA complex components RPA1 (70kDa subunit) and RPA3 (14kDa subunit) were not present in the original sgRNA library^46^, we targeted RPA1 and RPA3 by CRISPR-KO to determine if the other RPA subunits play a role in TAD boundary architecture, or if this RPA2 function was independent of the RPA complex. Indeed, deletion of each individual subunit increased *MYC* TAD boundary distances (Fig. 2B-C). Knockout of RPA1 or RPA2 increased TAD boundary distances from a mean of ∼400 nm in controls to ∼750 nm (p-value: 1x10^-8^) and ∼900 nm (p-value: 1x10^-8^), respectively, while RPA3 knockout led to a more moderate separation of ∼550 nm (p-value: 3x10^-7^) (Fig. 2B). Similar results were obtained by siRNA depletion (Extended Data Table 2). These observations indicate that the RPA complex as a whole is required for TAD boundary integrity.

**Fig. 2.**
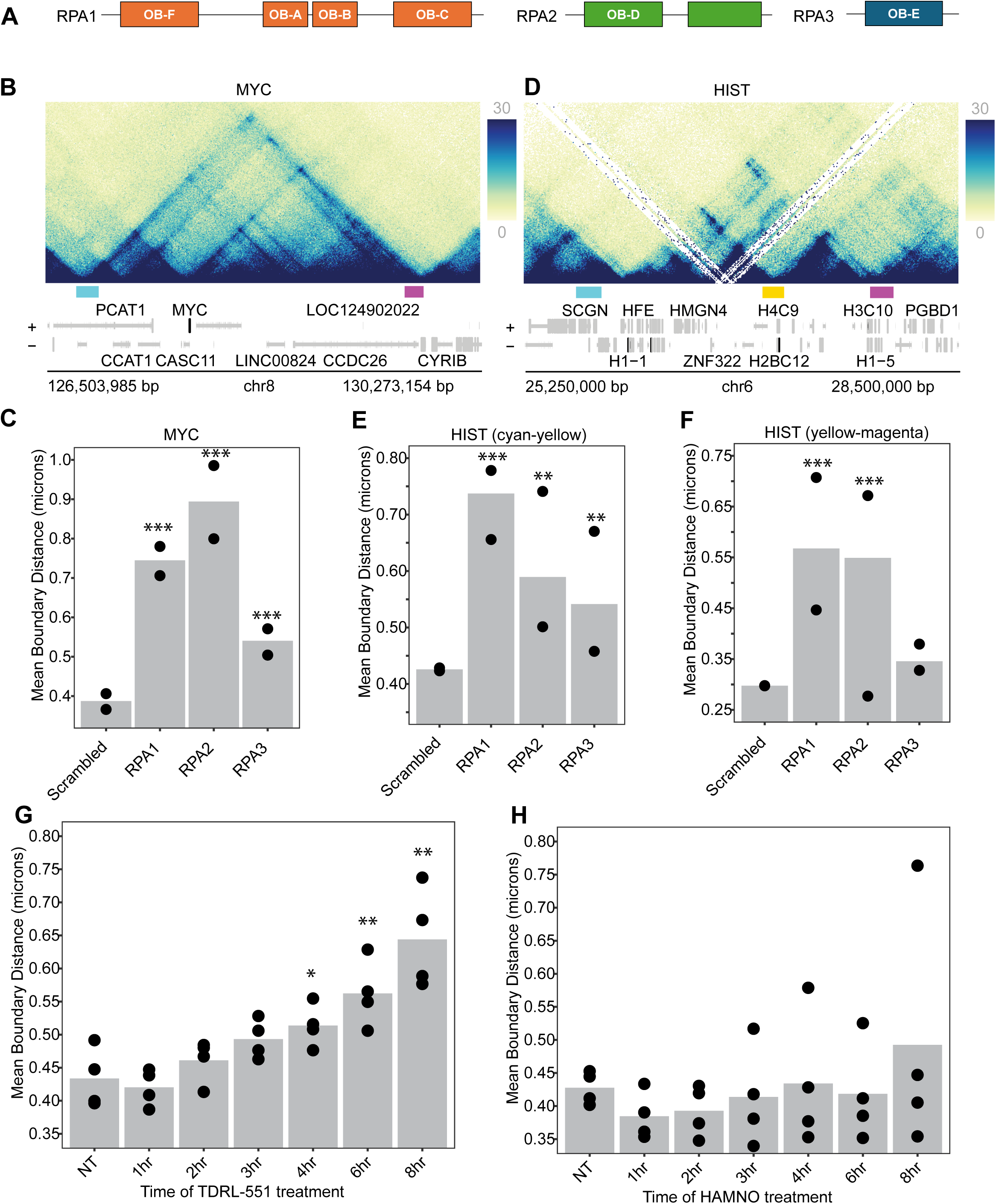
The RPA complex modulates TAD boundary structure. **A)** Schematic representation of the subunits of the heterotrimeric RPA complex with oligonucleotide/oligosaccharide binding (OB) fold DNA binding domains depicted for each subunit. **B)** Diagram of the *MYC* TAD. TAD FISH probe locations are indicated (cyan = 5’ probe, magenta = 3’ probe). **C)** Average TAD boundary distance (bars) after CRISPR-KO of RPA subunits at the *MYC* TAD from two technical replicates (black dots). Statistical analysis performed by ANOVA-Dunnett’s test, *** indicates p-value <0.001. **D)** Diagram of the *HIST* TAD and two sub-TADs. Location of FISH probes are indicated by rectangles for 5’ TAD boundary (cyan), 3’TAD boundary (magenta), sub-TAD boundary (yellow). **E-F)** Average TAD boundary distance (bars) after CRISPR-KO of RPA subunits at the *Hist* TAD from two technical replicates (black dots). Statistical analysis performed by ANOVA-Dunnett’s test, **P <0.01, ***P <0.001. **G-H)** Average *MYC* TAD boundary distances (bars) after RPA inhibition by TDRL-551 or HAMNO for indicated period of time. Dots represent individual biological replicates. Statistical analysis performed by two-tailed t-tests; *P< 0.05, **P< 0.01. At least 500 cells were analyzed per sample.

We next determined whether the role for the RPA complex in TAD structure was limited to the *MYC* TAD. Therefore, we analyzed the effect of loss of the RPA subunits on the histone (*HIST*) TAD located on human chromosome 6, which is smaller than the *MYC* TAD with two sub-domains of ∼1 Mb each (Fig. 2D). We labeled the sub-TAD boundaries at the *HIST* TAD after RPA knockdown and found that *HIST* TAD boundary distances at both sub-domains increased significantly after RPA1 deletion compared to scrambled cells similar of the behavior of the *MYC* TAD (Fig. 2E-F). These data indicate that the RPA complex is generally involved in boundary architecture at TADs (see also below).

Each RPA subunit makes direct contact with DNA via oligonucleotide/oligosaccharide binding (OB) folds^46^ (Fig. 2A). The small molecule TDRL-551 targets the ssDNA binding domain of RPA1 and is a potent inhibitor of RPA-DNA interactions^46–49^. Consistent with sgRNA and RNAi knockdown of RPA subunits, treatment of asynchronously growing HCT116 cells with TDRL-551 over the course of 8 hours resulted in a gradual increase with time in boundary distances at the *MYC* TAD from ∼450 nm in control cells to ∼650 nm (Fig. 2G; p value: < 0.005). Since the RPA complex is also involved in the DNA damage response (DDR) pathway by binding to damaged DNA and signaling for repair by ATR and RAD51^50^, we asked if RPA regulation of TAD boundary proximity worked indirectly through the DDR via ATR. We used the inhibitor HAMNO, which binds to the OB-F domain of RPA1, blocking ATR autophosphorylation and ATR phosphorylation of RPA2 at serine 33^51–54^. Treatment of asynchronous cells with HAMNO over the course of 8 hours did not cause discernable changes in boundary distance (Fig. 2H). Furthermore, RPA inhibition did not induce γH2AX (Extended Data Fig. 2C) excluding the possibility that RPA affects TADs boundaries via secondary DNA damage. These data suggest that the replication-associated RPA complex is directly involved in organizing TAD boundary architecture.

### TAD boundary dynamics during replication

The chromatin landscape is tightly regulated at the replication fork where chromatin features such as loops and epigenetic marks must be re-established after DNA synthesis^55,56^. Notably, some origins of replication initiate at TAD boundaries^57–60^. The identification of the RPA complex as a prominent determinant of TAD architecture prompted us to first characterize the behavior of TAD boundaries pairing during the cell cycle, particularly in S-phase. We mapped the behavior of the *MYC* TAD boundary at different phases of the cell cycle by performing DNA FISH followed by EdU labeling in asynchronous cells. The intensity of DAPI staining combined with EdU labeling was used to stage each cell in the cell cycle as previously described (Extended Data Fig 3A-B)^40^. Measuring the distance between the *MYC* TAD boundaries in each cell cycle phase, we observed that S-phase cells exhibited a moderate, yet consistent, increase in the distribution of boundary distances with averages of 410 nm compared to 367 nm and 321 nm in G1- or G2/M cells, respectively (Extended Data Fig 3C; p value = 0.043 and 0.088 respectively). These results point to conformational changes in TAD boundary architecture during S phase.

To more precisely characterize the behavior of the *MYC* TAD boundary during replication, we synchronized cells at the G1/S boundary by standard double thymidine block (see Methods) and measured TAD boundary distances after release into S-phase for various periods of time up to 8 hours. Based on RepliSeq mapping, TADs replicate as organized units, often initiating at the boundaries with the *MYC* TAD replicating during early-S-phase followed by replication bubbles converging in the middle of the TAD^17^ (Fig. 3A). We observe MYC TAD boundary separation concomitant with replication. Boundary distances increased from ∼380 nm in asynchronous cells to ∼500 nm in G1/S arrested cells (Fig. 3A; p-value: 0.022). As cells progressed through S-phase and TADs replicated, boundary distances gradually decreased back to ∼400 nm by the time cells reached G2/M-phase, 8 hours after release (Fig. 3A). These results are consistent with transient unfolding of TADs during replication.

**Fig. 3.**
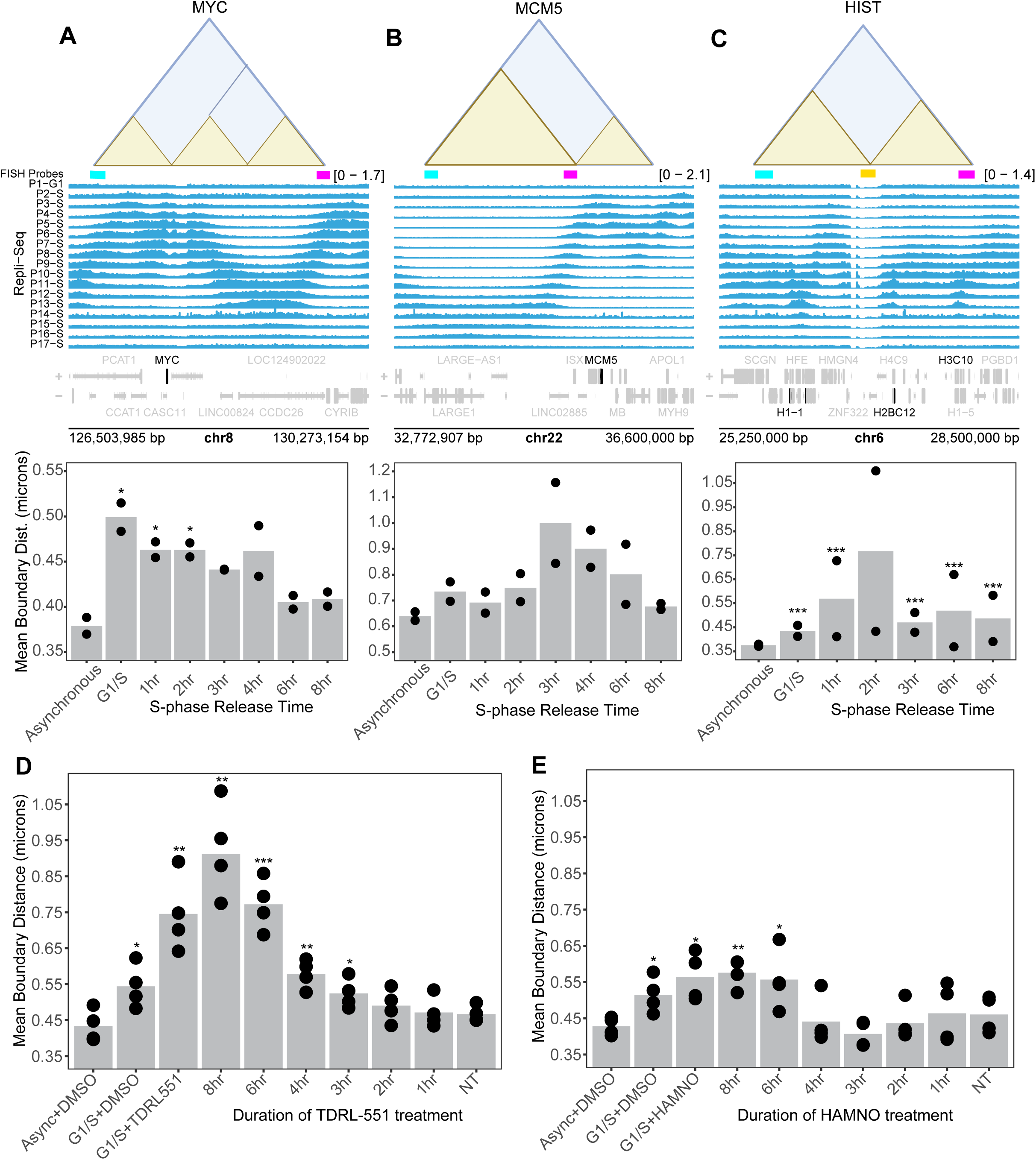
Inhibition of the RPA complex interrupts the memory of TAD boundaries after replication. **A-C)** Repli-seq profile from HCT116 cells in G1-phase and seventeen sequential time points during S-phase of the cell cycle as previously reported^17^. Early-, mid/nonspecific-, and late-replication timing regions of the genome are represented by the (**A**) *MYC*, (**B**) *MCM5* and (**C**) *HIST* TADs. Average TAD boundary distances (bars) after release for indicated period of time from G1/S arrest were measured at each TAD. Dots represent individual technical replicates. Statistical analysis performed by ANOVA-Dunnett’s t-tests, *P <0.05, ***P <0.001. **D-E)** Average TAD boundary distance measurements (bars) after (**D**) TDRL-551 inhibition or (**E**) HAMNO inhibition of RPA in S-phase synchronized cells. Dots represent individual biological replicates. Statistical analysis performed by two-tailed t-tests; *P< 0.05, **P< 0.01, ***P< 0.001. At least 250 cells were analyzed per sample.

We hypothesized that the observed separation of *MYC* TAD boundaries likely reflects the conformational changes the TAD undergoes as it replicates, returning to normal boundary architecture after completion of its replication. A prediction from this hypothesis is that TAD boundaries of late replicating TADs should separate later in S-phase. To test this prediction, we analyzed the late replicating ∼2.4 Mb *MCM5* TAD (Fig. 3B)^17^,as well as the *HIST* TAD, which exhibits a largely homogenous and unsynchronized pattern of replication during S-phase (Fig. 3C)^17^. Indeed, we found a clear relationship between replication timing and boundary separation for these TADs. In line with its later timing, maximum separation of the *MCM5* TAD boundary was only reached at 3-4 hr post-release into S-phase (Fig. 3B), whereas TAD boundary separation occurred throughout S-phase at the *HIST* 5’sub-TAD (Fig. 3C). These results suggest that TAD boundaries separate and re-establish proximity according to their replication timing during S-phase. The observed behavior is in line with the notion that TADs are highly dynamic structures^7^.

### Inhibition of RPA-DNA interactions interferes with re-establishment of TAD boundaries during replication

Considering that RPA affects TAD boundary distances as well as its known role in S-phase, we next wanted to determine whether RPA-DNA interactions were important for re-establishing TAD boundary proximity in a replication-dependent manner. When we treated asynchronous cells with the RPA inhibitor TDRL-551 for 1-8 hours and measured TAD boundary distances, we observed increased TAD boundary distances at each stage of the cell cycle (Extended Data Fig. 4A), consistent with a role of RPA in TAD replication. Single-cell analysis of cell-cycle staged cells shows that this effect is due to the larger proportion of S-phase cells, which have separated boundaries, with increasing treatment time (Extended Figure 3D, 4A).

To probe the effect of RPA on TAD boundary behavior during S-phase more directly, we synchronized cells at the G1/S-boundary and treated them with inhibitor at multiple time-points upon release (Fig. 3D) (see Methods). Progression of TDRL-551 treated cells through S-phase was only moderately reduced compared to untreated cells excluding the possibility that the observed effects are due to delayed replication (Extended Data Fig. 3E, 4B). TDRL-551 treatment increased *MYC* TAD boundary distances from ∼450 nm in asynchronous cells and ∼550 nm in G1/S arrested cells to ∼750 nm in TDRL-551 treated cells at the G1/S boundary (Fig. 3D, Extended Data Fig. 4B, p-value= 0.002). The effect was maximal in cells exposed to TDRL551 for the entire duration of S-phase, but importantly, no significant effect on *MYC* TAD boundary distances was observed when RPA treatment was initiated after *MYC* replication was complete at ∼4 hours post-release into S-phase (Fig. 3D). We also observed a similar, but more modest trend for HAMNO treated cells (Fig. 3E). These data demonstrate that the RPA complex plays a role in replication timing-dependent conformational re-organizations events of TAD boundaries during S-phase.

### Disrupting the RPA complex affects RAD21 loading

Our results suggest RPA binding to DNA is essential for organizing TAD boundary architecture across the cell cycle. To understand the mechanistic basis of this effect and given the prominent role of the cohesin complex in TAD formation, we asked if RPA-DNA interactions are important for cohesin association with chromatin. We performed ChIP-qPCR targeting RAD21 and CTCF at the *MYC* TAD in asynchronously growing cells in the presence or absence of RPA inhibitor TDRL-551. We observed a consistent reduction of RAD21 and CTCF binding upon RPA inhibition, albeit with distinct patterns depending on the location in the TAD. The two closely located RAD21 and CTCF binding sites 14.3 kb apart at the very 5’ *MYC* TAD boundary had different responses to RPA inhibition by TDRL-551. The upstream binding site 1 showed no change in CTCF nor RAD21 enrichment after RPA inhibition (Fig. 4B-C) whereas at binding site 2, RAD21 and CTCF enrichment decreased by 45% and 40% respectively (Fig. 4B-C). RAD21 enrichment at binding site 3 at the second sub-TAD boundary and binding site 4 located within the third sub-TAD decreased by 40% and 35%, respectively (Fig. 4 B-C). Although we observed a 50% decrease in CTCF enrichment at binding site 3, change of CTCF binding at site 4 was insignificant (Fig. 4 B-C). Interestingly, RPA inhibition led to the strongest decrease in RAD21 and CTCF enrichment at binding site 5 located at the 3’TAD boundary by 50% and 65%, respectively (Fig. 4 B-C). These data suggest that the RPA-DNA interactions are important for the association of RAD21 and CTCF with chromatin in asynchronously growing cells.

**Fig. 4.**
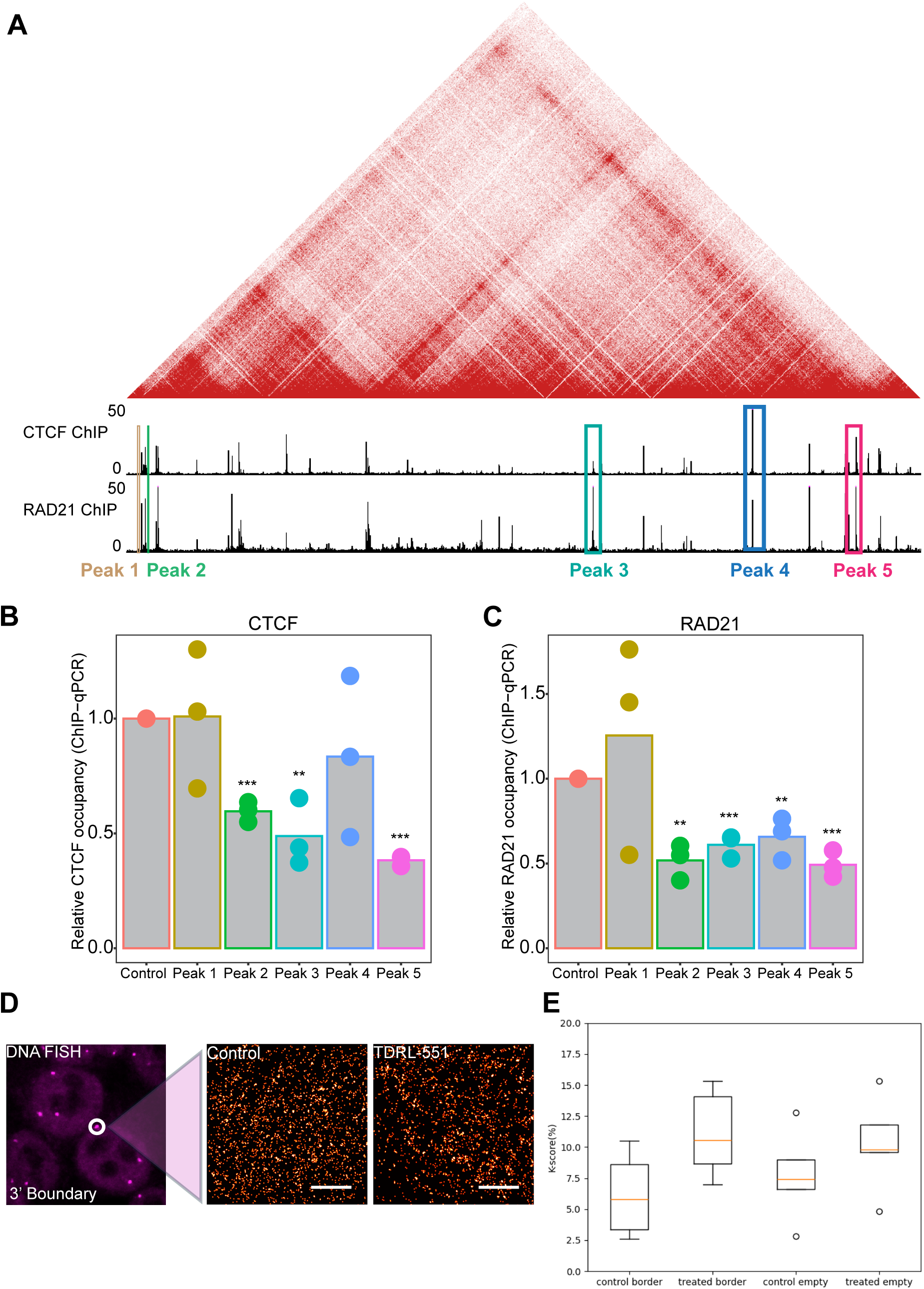
Inhibiting the RPA complex effects RAD21 loading and spatial distribution in the nucleus. **A**, HiC contact map with ChIP-seq tracks for CTCF and RAD21 at the *MYC* TAD in HCT116 cells^36,75^. **B-C)** ChIP-QPCR for **B)** CTCF and **C)** RAD21 enrichment across the *MYC* TAD including at the 5’ and 3’ *MYC* TAD boundaries in asynchronous cells treated with TDRL-551 for 6 hours. Values indicate averages from individual biological replicates (dots); Colors represent individual RAD21/CTCF peaks indicated in (**A).** Statistical analysis performed by two-tailed t-tests; *P< 0.05, **P< 0.01, ***P< 0.001. **D)** Single-molecule MINFLUX imaging of RAD21 at the *3’MYC* TAD boundary (dashed line) in asynchronous cells treated with TDRL-551 for 5 hours. DNA-FISH labeling of TAD boundaries was used to detect the field of imaging and labeling of RAD21 was performed by DNA-PAINT. Individual RAD21 molecules within the TAD boundary identified by DNA-FISH were imaged by MINFLUX nano-resolution microscopy. Signals represent individual molecules of RAD21. Scale bar: 500 nm; FISH-DNA panel (left), MINFLUX panels: 2x2µm. **E)** Distribution of RAD21 at *MYC* TAD boundaries and in arbitrary regions of the nucleus each in 2x2µm field of view. Clustering was measured by Ripley’s K-score. Values represent medians (red lines), 25 and 75 percentiles (box). 4-6 cells per measurement.

As an alternative approach to probe the association of cohesin with TAD boundaries we used single molecule visualization using MinFLUX (Minimal fluorescence photon fluxes) Imaging^61^. MinFLUX is an ultra-sensitive method for quantitative detection of single molecules with nanometer resolution in intact cells^62^. We visualized the 3’ MYC TAD boundary by DNA-FISH and measured the distribution of endogenous RAD21 by the immunolabeling approach DNA-PAINT^63^ at the TAD boundary in the presence or absence of TDRL-551 (Fig. 4D) (See Methods). Qualitative inspection indicated a change in spatial distribution of single RAD21 molecules from a largely homogenous distribution to a more clustered pattern with fewer individual signals at the 3’ TAD boundary in TDRL-551 treated cells compared to control cells (Fig 4D; Ext. Data Fig 5A). The qualitative assessment was confirmed by quantitative measurement of the heterogeneity in local molecular densities of RAD21 protein distribution using Ripley’s K-function (K-score)^64^ and nearest neighbor distance (NND) distributions^65,66^ (see Methods). We find larger K-scores indicating more heterogeneity in RAD21 protein densities in TDRL-551 treated cells compared to DMSO control cells. Furthermore, NND distributions from TDRL-551 treated cells exhibit more deviation from a uniform random distribution (Fig 4E; Ext. Data Fig 5B). Similar changes in RAD21 localization occurred in random regions outside of the *MYC* TAD boundaries, likely reflecting cohesin binding sites throughout the genome (Fig. 4E). These results suggest reduced binding of RAD21 upon inhibition of RPA, in line with a role of RPA in recruitment of the cohesin complex to chromatin.

## Discussion

We have performed a high-throughput imaging CRISPR knockout screen to identify effectors of TAD architecture. Using TAD boundary distance as a measure of TAD organization, we find that the RPA complex is critical in ensuring faithful re-establishment of TADs after replication. Our results highlight the role of cell cycle regulators and replication factors in organization of chromatin domains.

We developed a DNA-FISH based high-throughput assay for the quantitative measurement of TAD boundary distances at individual alleles and used it in a CRISPR screen of more than 1000 nuclear proteins^39–41,64^. Reassuringly, the top hits included multiple subunits of the cohesin complex including RAD21, NIPBL, and MAU2, validating our screening approach. Importantly, we identified several novel groups of modulators of TAD architecture, including cell-cycle proteins and replication factors. One of the most prominent hits was RPA2, a component of the RPA complex involved in replication and recombination. Loss of the other RPA complex components RPA1 or 3 and inhibition of the RPA complex by small molecules phenocopied loss of RPA2 and caused disruption of TAD architecture as indicated by separation of the domain boundaries. These effects were not due to recombination or the DDR since loss of the DDR factors ATM and RAD51 did not phenocopy them. Although deletion and long-term inhibition of RPA causes aneuploidy of *MYC*, acute inhibition of RPA on the order of hours does not cause DNA damage nor aneuploidy, ruling out secondary effects.

The identification of the RPA complex, involved in replication, points to an important role of S-phase progression in TAD structure. Indeed, in time-course experiments we found a correlation between replication timing and separation of TAD boundaries. Specifically, we noticed separation of TAD boundaries shortly prior to the time of replication. For example, TAD boundaries of the early replicating *MYC* TAD were already separated in G1/S-arrested cells, whereas the boundaries of the late replicating *MCM5* TAD were still proximal to each other in G1/S arrested cells and only separated 3hrs into S-phase just prior to replication of the *MCM5* TAD around 4-6h into S-phase. These results suggest that separation of TAD boundaries is an initial step in TAD replication and that boundary pairing and TAD architecture is re-established upon completion of replication. Loss of RPA complex function interferes with this process, leading to prolonged separation of TAD boundaries.

These findings are in line with Repli-seq data, which show that TADs tend to replicate as a unit with specified replication initiation zones^14,60^. Furthermore, TAD replication begins from the boundary elements and proceeds inward towards the middle of the TAD until replication origins converge^57,58^. Our data suggest that TAD boundaries separate and rejoin in a replication timing-dependent way. We suggest that the separation of TAD boundaries reflects relaxation of the TAD structure ahead of the replication fork initiation, causing TAD boundaries to move apart, likely to facilitate access of the replication machinery. For early replicating TADs like *MYC*, boundaries may separate at the G1/S checkpoint before proceeding into S-phase and considering that origin recognition complexes associate with chromatin throughout the cell cycle^67^, it is possible that separation of boundaries at early replicating TADs may be required for entry into S-phase. We find that CRISPR-KO of RPA components has similar effects on TAD boundary separation as knockout of the cohesin component RAD21. Furthermore, short-term TDRL-551 inhibition of RPA and acute depletion of RAD21 resulted in the same phenotype. These similarities point to an interplay between RPA and cohesin in establishing and maintaining TAD boundary architecture. In support, in vitro assays performed with tethered circularized DNA and purified cohesin and RPA show that RPA binding to DNA is required for proper loading of cohesin onto single stranded DNA, similar to association at the replication fork^68^. Furthermore, RPA, along with the replicative cohesin subunit CDCA5, is important for chromatid cohesion during S-phase and interacts with the MCM replication machinery^27^. In line with a mechanism by which the RPA complex facilitates cohesin recruitment to TAD boundaries, we demonstrate that RPA inhibition leads to reduction of RAD21 association with chromatin at the *MYC* TAD boundary. This reduction coincides with a change in nuclear distribution of RAD21 at the single molecule level as demonstrated by ultra-high resolution imaging. We also find altered CTCF binding upon RPA inhibition, suggesting that in the absence of RPA function, CTCF recruitment to chromatin is disrupted, which may lead to altered loop extrusion by cohesin, and possibly changes in TAD architecture. Alternatively, the inability of cohesin to bind to boundaries in the absence of RPA may be due to alterations in chromatin structure which lower the affinity of cohesin to chromatin. Further biochemical and cell-based assays will be required to delineate the details of the molecular interplay between RPA and cohesin complexes.

Taken together our results have identified replication factors as prominent modifiers of TAD architecture during interphase. Our findings indicate that TAD structure is temporarily lost during S-phase and must be faithfully re-established upon completion of replication and that close interplay between the RPA replication complex and the cohesin machinery is critical for this process. Our study opens the door to begin to understand in depth the relationship between cell cycle progression and TAD organization in time and the three-dimensional space of the nucleus.

## Acknowledgement

This work utilized the computational resources of the NIH HPC Biowulf cluster (https://hpc.nih.gov). HCT116-Cas9 clone A cells were kindly provided by Scott E. Martin at Genentech. This research was supported by the Intramural Research Program of the National Institutes of Health (NIH), National Cancer Institute NCI, Center for Cancer Research through grant 1-ZIA-BC010309 to TM and grant 1-ZIA-BC-011567 to HiTIF. The CCR/ LRBGE Optical Microscopy Core is funded by the Intramural Research Program of the National Cancer Institute (NCI), Center for Cancer Research (CCR) under project number 1-ZIA-BC-011574 to TK. TB was also supported by a NIGMS Postdoctoral Research Associate Training (PRAT) Program (1FI2GM146623-01), and a NCI Intramural Continuing Umbrella of Research Experiences (iCURE) Postdoctoral Fellowship. The contributions of the NIH author(s) were made as part of their official duties as NIH federal employees, are in compliance with agency policy requirements, and are considered Works of the United States Government. However, the findings and conclusions presented in this paper are those of the author(s) and do not necessarily reflect the views of the NIH or the U.S. Department of Health and Human Services.

## Authors’ contributions

TB, TM conceptualized the project, designed the experiments, and interpreted the results. TB, FA executed experiments. AK, MF performed computational imaging processing and analysis. KSMR contributed to HiC data analysis. LO contributed to high-throughput imaging experiments. TB and TM drafted the manuscript. All authors participated in reviewing and editing the manuscript. TM, GP, and TK obtained funding.

## Competing interests

The authors declare they have no competing interests.

## Methods

### Cell culture

Cells were cultured as previously described^42^. HCT116-Cas9^69^ cells were cultured in RPMI-1640 medium L-glutamine, HEPES, pyruvate, glucose, sodium bicarbonate (ATCC, 30-2001) supplemented with 10% fetal bovine serum (Gibco, 10-082-147), 1% penicillin-streptomycin (GIBCO, 15140-163), and Blasticidin (InvivoGen, ant-bl-05) at 37°C in 5% CO2. HCT116-RAD21-AID degron cells^70^ were cultured in McCoy’s 5A medium supplemented with 2 mM L-glutamine and 10% fetal bovine serum, and 1% penicillin-streptomycin.

### Cell Cycle Synchronization

To perform S-phase analysis, we synchronized cells by standard double thymidine treatments at the G1/S-phase checkpoint^40,64^. In brief, cells were treated with 2mM thymidine for 16 hours, then released into fresh growth medium without thymidine for 8 hours. The cells were again treated for 16 hours with 2mM thymidine, blocking them at the G1/S-phase checkpoint. Synchronized cells were release into S-phase for 0-8 hours before PFA fixation.

### RPA inhibition

HCT116-Cas9 cells were seeded in 96-well plates (30,000 cells/well) and grown for 72 hours before treatment. Duplicate wells were treated with RPA inhibitors at IC50 concentrations of 18µM for TDRL-551^46–49^ or 50µM for HAMNO^51–54^ for 1-8 hrs before PFA fixation. For S-phase assays, cells were synchronized at the G1/S boundary by double thymidine treatment. All blocked cells were released into S-phase for 8hrs by washing 1 time with PBS, 2 times with fresh growth media, and treated with RPA inhibitors for 1-8hours before PFA fixation. Asynchronous and S-phase cells were then processed for DNA HiFISH.

### HiFISH

High-throughput FISH was performed as described in^18,39,40^. Cells were seeded in Perkin-Elmer CellCarrier 384-well plates by multi-drop automation or manually by multi-channel pipette at 1,500 cells per well (37,500 cells/ml) and were allowed to grow for 72 hrs. A BlueWasher dispenser (BlueCatBio) was used to automate cell processing before probe hybridization. Cells were fixed in 4% paraformaldehyde in PBS for 10 min, washed in PBS three times, permeabilized in Triton X-100 with saponin in PBS for 15 min, washed two times in PBS, incubated in 0.1N HCL for 12 min, washed in 2X SSC for 5min, then incubated at 4°C overnight in 50% formamide/2x SSC. Nick translated BAC probes in hybridization mixture were dispensed into each well using a Mosquito HV Genomics liquid handler (TTP Labtech). Hybridization was carried out by denaturing probes and cellular DNA together at 85°C for 8 min then incubated at 37°C for 72 hours. Excess probe was removed from wells by three washes of 2X SSC and three washes of 1X SSC using the BlueWasher. The DNA was stained by 5 ng/ul 4′,6-diamidino-2-phenylindole (DAPI) in PBS for 5 min then washed and stored in PBS before imaging.

### Probe generation

DNA FISH probes were designed to label TAD boundary regions as follows. Genomic regions were identified based on publicly available HiC sequencing heat maps^36^. We selected TAD boundaries by the presence of corner dots indicating contacts between boundary elements. To generate probes, we chose from the UCSC genome browser multiple BACs (Chori/BACPAC) that tiled the boundary elements. Single BAC clones were picked from colonies streaked onto chloramphenicol agar plates. BAC plasmids were isolated using the NucleoBond BAC 100 kit (Macherey-Nagel, cat # 740579). Fluorescent labeling with Dyomics (Germany) DY-488 Conjugated dUTP (488-34), Dyomics DY-549P1-dUTP (549P1-34), or DY-647P1-dUTP(647P1-34), was performed as previously described by nick translation and precipitation protocols^18,39,40^.

Final selection of boundary probes was by empirically testing and optimization for each TAD by performing DNA FISH. Probe coordinates are provided in Extended Data Table 3.

### CRISPR-KO Library Design

A previously described custom-made arrayed synthetic sgRNA library targeting 1,064 genes associated with chromatin biology and nuclear architecture was used^64^. The sgRNA library was reformatted to 384-well plates using a PerkinElmer Janus and a Beckman Coulter ECHO525 liquid handlers at a final concentration of 0.25 pmoles/μL. Each gene was targeted by 3 pooled sgRNA oligos in the same well.

### CRISPR-KO Screens

For reverse transfection in 384-well format, 325 nL of library sgRNA (0.25 pmoles/μL) was spotted into each well (0.08 pmoles/well) of Perkin-Elmer CellCarrier 384-well plates using an ECHO525 acoustic liquid handler (Beckman). Control wells containing 7 replicates each of non-targeting scrambled control sgRNA (Synthego, Cat. No. 063-1010-000-000) or sgRNAs targeting PLK1, NIPBL, or RAD21 were included on each plate. The PLK1, NIPBL, or RAD21 targeting sgRNAs were obtained as pooled gene knockout kits from Synthego (1.5 nmol). All control sgRNAs used identical chemistry and quantities as the library sgRNAs. Spotted plates were dried at room temperature under laminar flow, sealed, and stored at -30°C until transfection.

On the day of transfection, spotted sgRNA plates were thawed for 30 minutes, equilibrated to room temperature, and briefly centrifuged at 365xg. After removing the seal, 20 μL of prewarmed serum-free OptiMem medium (Thermo Fisher Scientific, Cat. No. 31985070) was dispensed into each well using a Multidrop dispenser (Thermo Fisher). The ECHO525 dispenser was used to add 50 nL of Lipofectamine RNAi MAX (Thermo Fisher Scientific, Cat. No. 13778075). Plates were incubated at room temperature for 30 minutes to allow RNA-lipofectamine complex formation. Subsequently, a cell suspension prepared in prewarmed OptiMem medium containing 20% FBS was dispensed using the Multidrop dispenser for a total volume of 40 μL and a final sgRNA concentration of 2 nm. Plates were incubated at room temperature in a laminar airflow hood for 30 minutes before transferring to a cell culture incubator for 72 hours. CRISPR-KO screens were performed in 3 biological replicates on separate days.

### Imaging

Fixed cells in 384-well plates were stained with DNA FISH probes and imaged using Yokogawa CV7000 or CV8000 spinning disk confocal microscopes following previously described parameters^42^. Samples were excited using 405 nm (DAPI), 488 nm (green fluorophores), 561 nm (red fluorophores), and 640 nm (far-red fluorophores) laser excitation lines combined through a 405/488/561/640 nm quad-band dichroic mirror. The emitted fluorescence was captured through a 60X water-immersion objective (NA = 1.2) and filtered through 445/45 nm (DAPI), 525/50 nm (green), 600/37 nm (red), or 676/29 nm (far-red) band-pass filters. Two 16-bit sCMOS cameras (2048 × 2048 pixels, 1 × 1 binning; effective pixel size = 0.108 μm) acquired image Z-stacks with 1 μm steps using real-time maximal projection. Nine fields per well were imaged across different experiments to acquire sufficient cell images.

### Image Analysis

Image analysis on Z-stack image projections was performed as previously described using HiTIPS^42^ . HiTIPS analysis settings were empirically optimized for typical nuclear diameter and for DNA FISH foci intensity/size characteristics. Nuclear segmentation utilized the GPU-accelerated CellPose algorithm^71^ and DNA FISH foci spot detection employed a Laplacian-of-Gaussian approach. Final spot coordinates were defined as the centroid of each segmented focus. Average nuclear fluorescence intensity for DAPI (405 nm) and EdU (640 nm) channels was measured at the single-cell level.

### Identification of Screen Hits

Single-cell and single DNA FISH spot data was exported as CSV files from HiTIPS. R 4.3.3^72^ was then used to filter cells that had at least 2 spots in each of the DNA FISH images and had the same number of spots in both the DNA FISH images. Euclidean distances between all possible DNA FISH spot pairs in the 2 channels were calculated and only the minimum value per red spot, per cell was retained for analysis. Spot-spot distances were then averaged on a per well basis and analyzed using the cellHTS2 package^73^ Per-well raw measurements were normalized per plate using the median value of sgRNA library treatments and the B-score method. All per-plate normalized values from a single biological replicate were further standardized using the Z-score. Z-score values for identical well and plate combinations across biological replicates were averaged to obtain Mean Z-scores. Screen hits were identified as genes whose knockout resulted in a mean Z-score above 2.5 for mean distance between green and red DNA FISH spots representing *MYC* TAD boundaries.

### MINFLUX

MINFLUX uses repeat scanning of the field using a doughnut-shaped laser beam. When a molecule is detected, the system locks on it and keeps repeatedly localizing the same molecule until the “imager” oligo detaches and fluorescence from the molecule ceases. This results in a trace of localizations associated with the molecule which we simply call trace. After that, MINFLUX continues to scan the field until the next molecule is detected. This repeat scanning of the field can lead to multiple localization traces from the same molecule in the final MINFLUX data^61,62^.

To obtain localization of single molecules by MINFLUX microscopy^61^, RAD21 was immunolabeled by the DNA-PAINT method^63,74^. In brief, cells grown on 18 mm coverslips for 72hrs were treated with TDRL-551 or DMSO for 6hrs. DNA FISH was carried out as described above to label the 3’ *MYC* TAD boundary in order to identify a region of interest by confocal microscopy. Immunolabeling of RAD21 was next carried out by the immunofluorescence protocol described above using monoclonal antibody Anti-RAD21 [Abcam, ab217678] followed by labeling with the DNA PAINT Massive-sdAB 2-Plex kit (Massive Photonics GmbH, Germany) containing anti-mouse camelid nanobody Massive-Anti-Mouse kLC Docker Strand 1, according to the manufacturer’s protocol. During this process, each primary antibody is bound by two identical secondary nanobodies providing a 2:1 labeling ratio. Coverslips with labeled cells were mounted into a LCI chamlide magnetic chamber (Quorum Technologies Inc., Canada) and covered with 0.5 ml of a proprietary MP imaging buffer from the kit. 500 pM of the MP Imager Strand 1 conjugated to the Atto655 fluorophore was added to the chamber to hybridize to the kLC Docker Strand 1. Multiple 2x2 μm fields of view were imaged on the MINFLUX imaging system (Abberior GmbH, Germany) equipped with an Olympus Objective UPlanSApo 100x/1.40 Oil 8/0.17/FN26.5. For each image, localizations of the individual molecules were collected in 2D for 6 hours, with 0.83 pinhole size and 12% of the 642 nm laser power.

To discard outliers, traces containing only one localization were removed. The trace average and standard deviation were calculated and reported as localization events. However, in the case of multiple localization events associated with a molecule, we combined localizations that are closer than the average localization precision (approximately 5 nm) to attain a single location with high precision for each molecule. This data is collected as a spatial map of individual molecules. Finally, we used the obtained molecule locations to find the spatial variation in molecule density by calculating the Ripley’s K-function and NND distributions^41,42,64–66^.

### siRNA

The siRNA validation library was generated based on the top 10 ranked z-score hits as well as 38 targets that ranked higher than CTCF for TAD boundary distance with z-scores over 1.27 (Ext. Data Fig. 2A). siRNAs were spotted at random in 384-well plates at 150 nl of 5 µM siRNA per well (0.75pmole/well), using siNIPBL and siRAD21 as positive controls. We performed the same automation pipeline for reverse transfection used for CRISPR deletion screen (see above). Single siRNAs were spotted per well with three different siRNA targeted against each hit. Cells were incubated for 72hr depletion before PFA fixation and DNA-FISH processing.

### ChIP

Cells (18 million) were grown in T75 flasks for 72 hours before treatment for 6hrs with DMSO or 18µm TDRL-551 in growth medium. The cells were then briefly detached by trypLE (Gibco Cat.# 12604013), resuspended in PBS, cross linked in 1% formaldehyde, flash frozen in 15mL conical tubes on dry ice and ethanol, then stored at -80°C. Chromatin was isolated and purified according to the manufacturer’s instruction using the Pierce Magnetic ChIP Kit (Pierce^TM^ Magnetic ChIP Kit; Cat # 26157). Nuclei were isolated from thawed cells in the kit’s Membrane Extraction Buffer containing Halt protease/phosphatase inhibitors followed by DNA digestion with MNase (Thermo Scientific; 1862753). The nuclei were then sonicated in IP Dilution Buffer containing Halt to extract chromatin for 20 cycles of 30sec on/30sec off using a Diagenode Biorupter Plus. Each sample was then separated into 3 tubes for overnight immunoprecipitation with Normal Rabbit IgG (Thermo Scientific; 186224), Mouse Anti-RAD21 (Abcam; AB217678), or Mouse Anti-CTCF (Abcam; AB128873). 10% input was set aside from each tube prior to addition of antibody for later use as a control. Chromatin-antibody complexes were pulled down by magnetic A/G beads provided in the kit, then washed in the kit’s 1X IP Dilution buffer, eluted with the kit’s 1X Elution buffer, and reverse-crosslinked according to the protocol. Isolated chromatin was purified using DNA Clean-Up Columns provided in the kit. Enrichment of RAD21 and CTCF at the *MYC* TAD was measured by qRT-PCR amplification of genomic regions known to have co-localized RAD21 and CTCF ChIP peaks based on ChIP seq data (Primers listed in Table)^75^.

### Statistical Analysis

Two-tailed t-test was used to find the p-value for average difference in TAD boundary distance taken from well level distributions measured per condition and replicates. ANOVA test followed by Tukey’s post-hoc test for the differences between mean TAD boundary distances across cell cycle phases. ANOVA test followed by Dunnet post-hoc test for the difference between TAD boundary distances for RPA inhibition and S-phase time course.

### Data Availability

All R scripts used for image quantification, including calculations of TAD boundary distances from DNA FISH data are publicly available at : https://github.com/CBIIT/tad-boundary-replication-analysis. ImageJ-generated JPEG composites underlying the microscopy panels, along with the DNA FISH datasets used in this study, have been deposited in Figshare (https://figshare.com/s/10c46d79c5a1491a95f7).

**Extended Data Table. 1 CRISPR-KO screen TAD boundary distances.** GeneSymbol column lists the genes targeted in the sgRNA library. Targets are listed sorted by z-score in the second column. Boundary distances in µm are listed in the third column.

**Extended Data Table. 2 Validation of screen hits and the RPA complex.** An siRNA validation library targeting 48 genes including top hits from the screen with a z-score of >1.5. The library also included targets that led to TAD boundary distances greater than CTCF knockout in the screen as well as siRPA1 and siRPA3.

**Extended Data Table. 3 BAC Probe Locations.** List of genomic locations for bacterial artificial chromosomes (BACs) used in this study.

**Extended Data Table. 4 Sequencing tracks.** Genomic sequencing tracks for HiC sequencing, ChIP-seq, and Repli-seq data used in this study.

---

**Extended Data Fig. 1.**
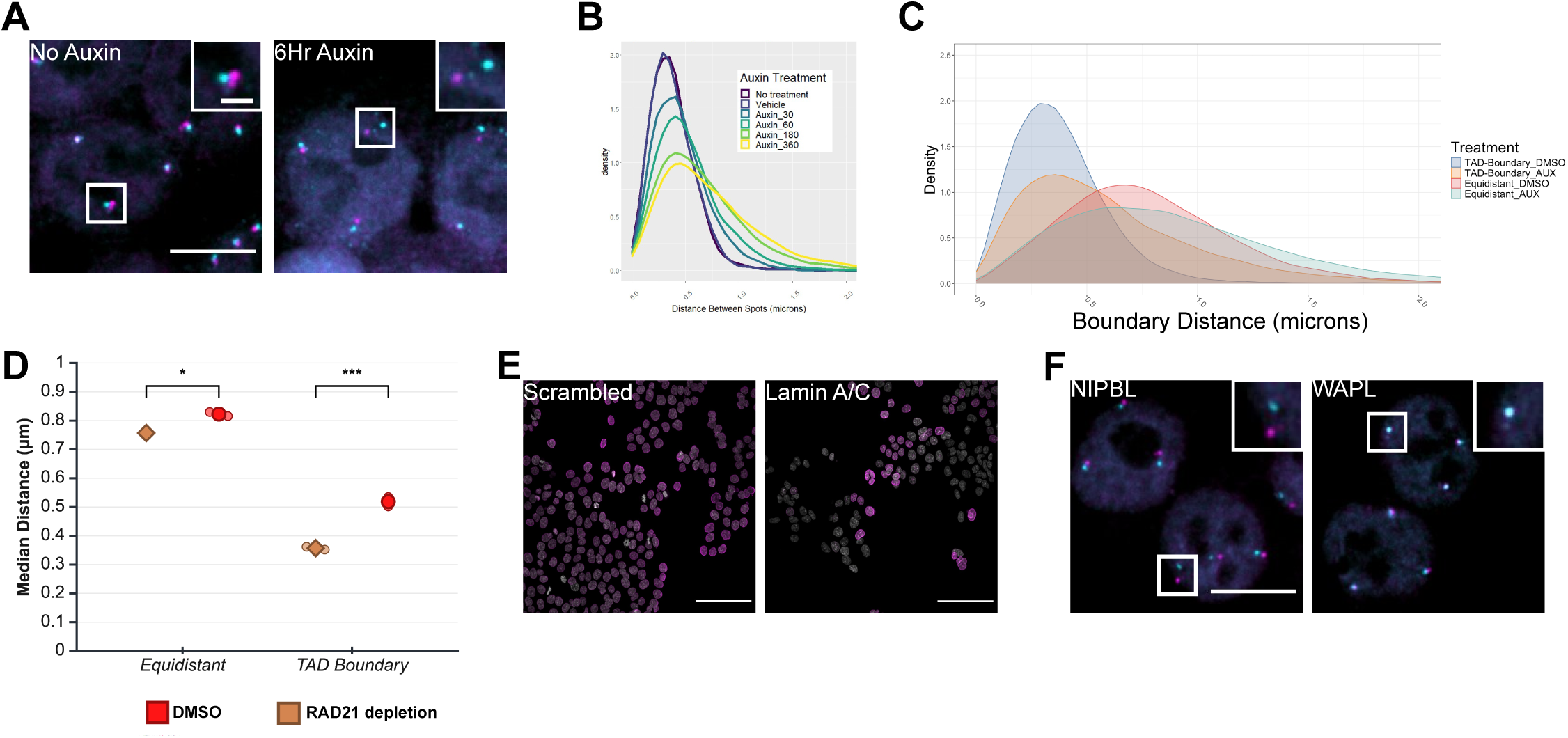
High-Throughput imaging pipeline to measure TAD boundary proximity. **A)**, HCT116-RAD21mAID cells were treated for 6 hours with auxin to deplete RAD21. BAC probes label the *MYC* TAD at the 5’ boundary (cyan, DY-488) and 3’ boundary (magenta, DY-549). scale bar: 10µm. **B)** Distribution of TAD boundary distances after degron-mediated depletion of RAD21 for 6 hours measured by HiTIPS automated image analysis. **C)** Distribution of MYC boundary distances (blue) and of the *MYC* 5’ TAD boundary relative to an equidistant non-boundary upstream region (red) in the presence of RAD21 (blue, red, respectively) or after RAD21 depletion for 6 hours (orange, light red, respectively). **D**) Median distances of distribution data shown in panel **C)**. Values represent means (large symbols) from three individual experiments (small symbols); More than 3,000 alleles were analyzed for each individual experiment. *p-value< 0.05, ***p-value< 0.001. **E)**, CRISPR-KO of LAMIN A and C in HCT116-Cas9 cells. Lamin A/C protein levels detected by immunofluorescence (magenta) and nuclei (DAPI, grey) were measured by Columbus software. Scale bar: 100µm. **F),** Representative images of the effect of knockout of architectural genes NIPBL and WAPL. 5’TAD boundary (cyan), 3’TAD boundary (magenta). Scale bar: 10µm.

**Extended Data Fig. 2.**
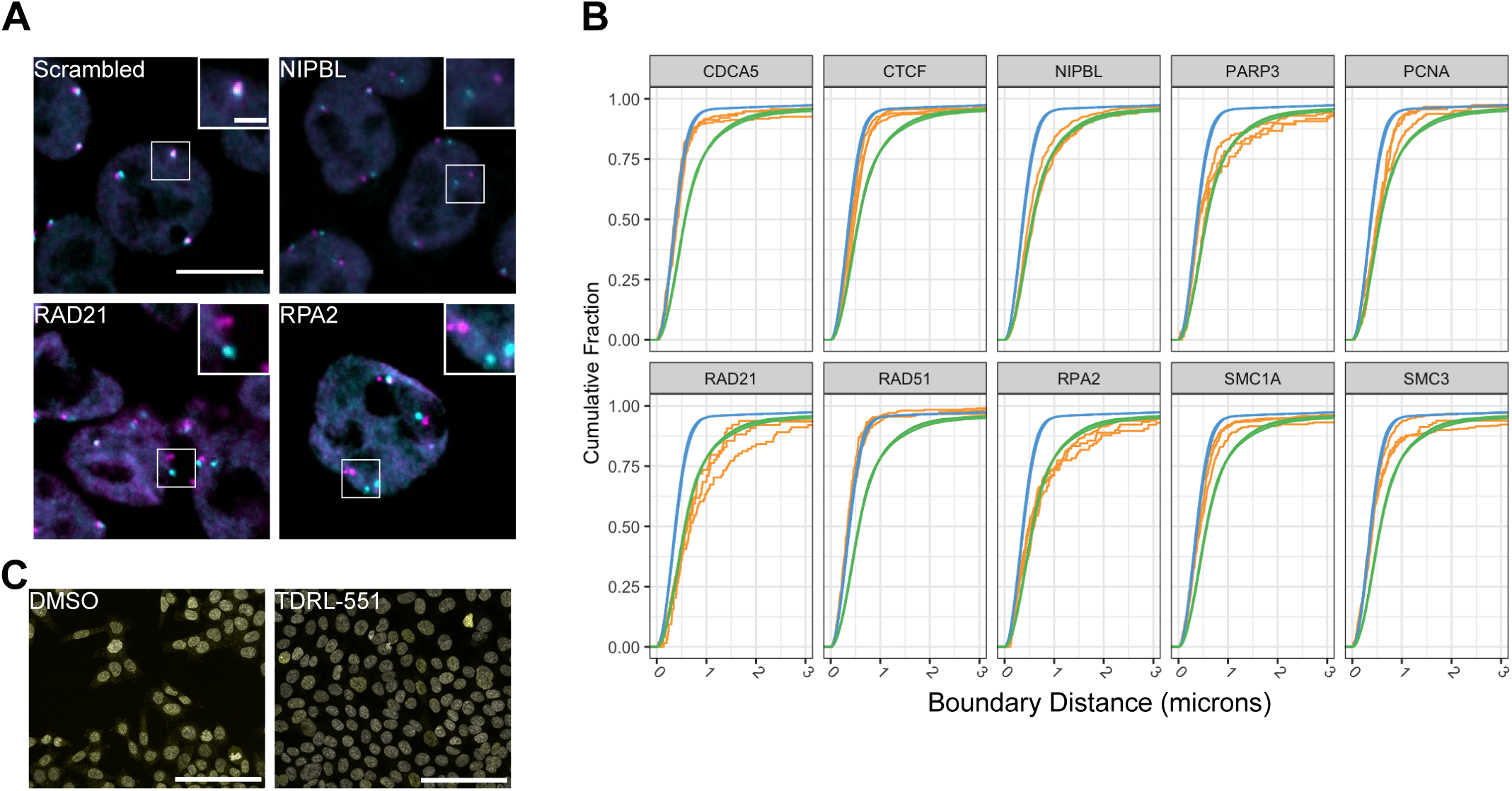
Validation of screen hits and the RPA complex. **A)** Representative images of siRNA-depleted cells for the RPA2 compared to controls NIPBL and RAD21. 5’ MYC TAD boundary (green), 3’ MYC TAD boundary (red), Scale bar: 10µm. **B)**, Cumulative allelic distribution of TAD boundary distances after CRISPR-KO screen. Curves represent targeted replication genes, cohesin subunits, and DNA damage response genes. Scrambled control (blue), NIPBL control (green), indicated target (yellow). Three biological replicates are shown. **C)**, Immunofluorescent labeling of γ-H2Ax in HCT116-Cas9 cells treated with DMSO or TDRL-551 for 6 hours. Nucleus (grey), γ-H2Ax (yellow), Scale bar: 100µm.

**Extended Data Fig 3.**
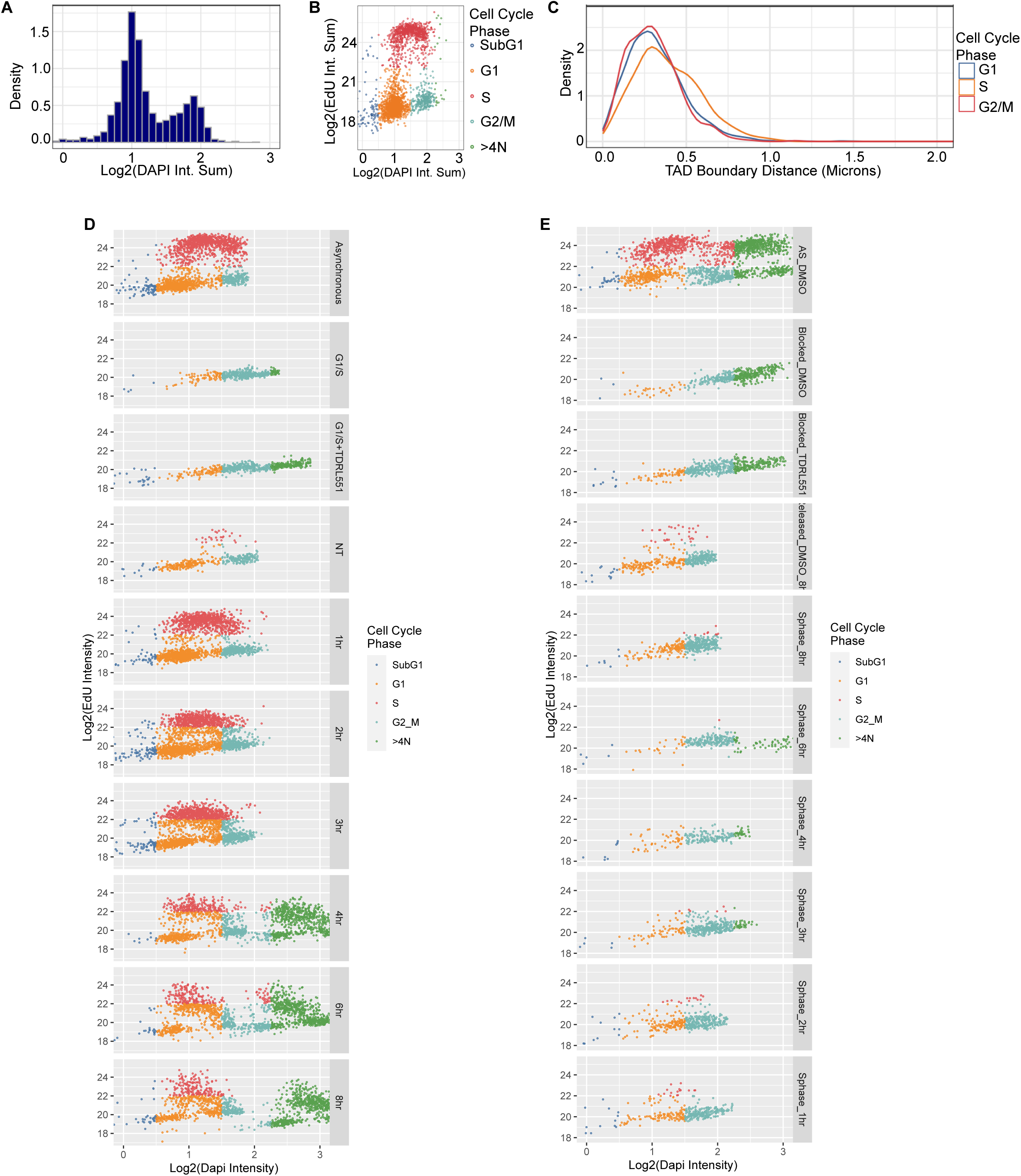
DAPI and EdU cell-cycle staging of cells. **A-C)**, Asynchronous cells were incubated with EdU for 1 hour before PFA fixation, DNA FISH, and Click reaction for EdU fluorescent labeling. **A)** DNA quantification by DAPI staining intensity. **B**) Assignment of cell-cycle phases based on DAPI intensity and Edu intensity. **C)** Allelic distribution of *MYC* TAD boundary distances in G1-, S-, and G2/M-phase staged cells. **D)** Cell-cycle analysis in asynchronous cells treated with RPA inhibitor TDRL-551 over the course of eight hours. **E)** Cell-cycle analysis in cells synchronized and released into S-phase treated with RPA inhibitor TDRL-551.

**Extended Data Fig 4.**
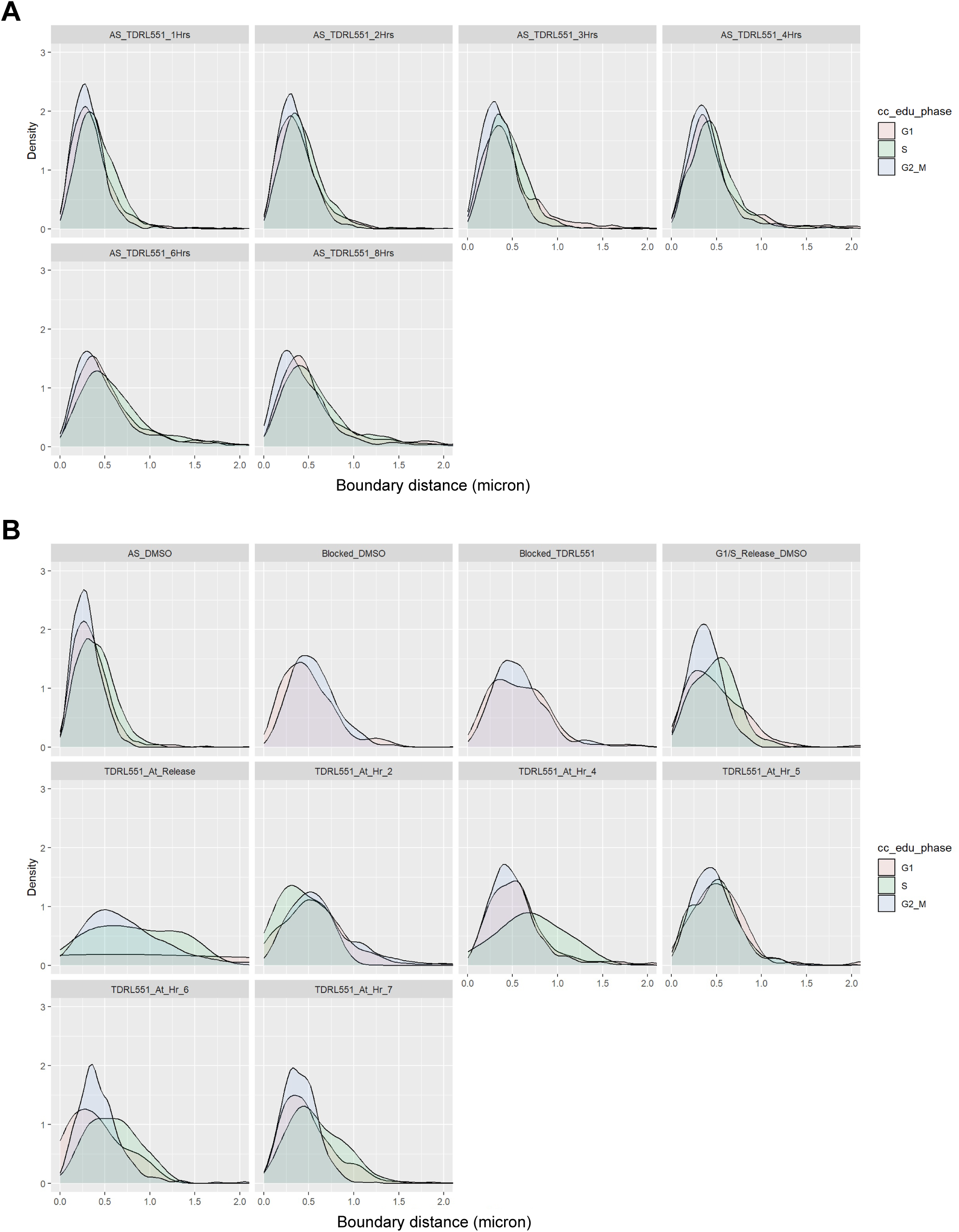
TAD boundary distance across the cell cycle. **A)** Asynchronous cells were treated with TDRL-551 for the indicated period of time and then incubated with EdU for 1 hour before processing for DNA FISH and EdU fluorescent labeling. Allelic distribution of *MYC* TAD boundary distances in G1-, S-, and G2/M-phase staged cells. **B)** S-phase synchronized cells were treated with TDRL-551 for the indicated period of time then incubated with EdU for 1 hour before processing for DNA FISH and EdU fluorescent labeling and allelic distribution of *MYC* TAD boundary distances was determined. Typically more than 1800 cells were analyzed per experiment.

**Extended Data Fig 5.**
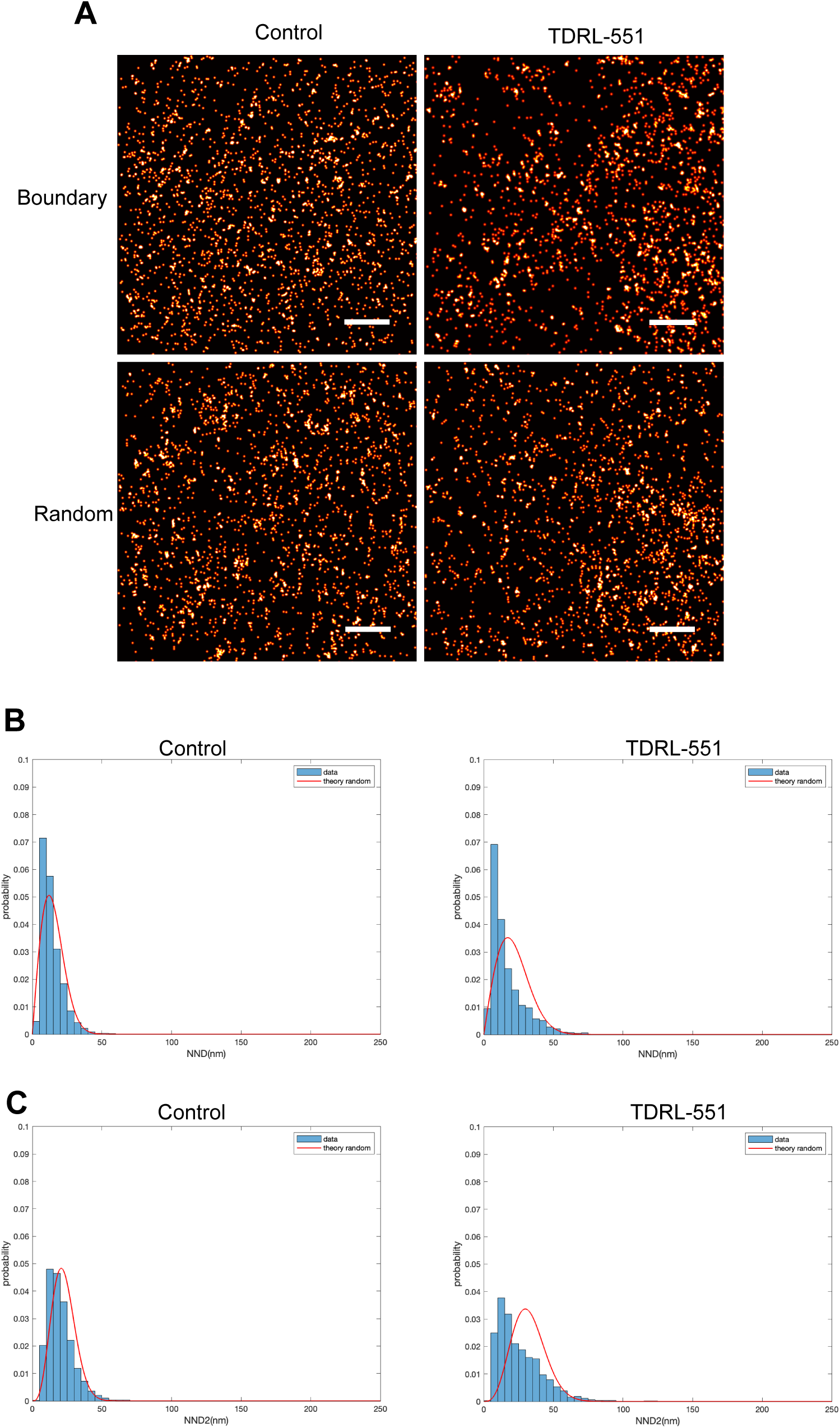
MINFLUX Nano-resolution microscopy of RAD21. **A)** Single molecule labeling of RAD21 by DNA-PAINT and imaged by MINFLUX nanoscale microscopy. Scale bar = 500 nm. Asynchronous cells were either not treated or treated for 6 hours with RPA inhibitor TDRL-551 and RAD21 visualized at the MYC TAD boundary identified by DNA FISH (boundary) or outside at a random location in the cell nucleus. **B-C)** Analysis of the nearest neighbor distance (NND) distributions between individual RAD21 molecules for the first nearest (**B**) and second nearest (**C**) neighboring molecule at the *MYC* TAD 3’ boundary in control cells compared to TDRL-551 treated cells. The red curves represent the expected NND distribution for uniformly randomly distributed molecules with densities similar to the imaged samples and the histogram represents the measured distribution of distances between single molecules of RAD21.

## Notes

### Competing Interest Statement

The authors have declared no competing interest.

### Summary of Updates

We have corrected the name of one of the co-authors (GP).

## References

1. Misteli, T. The Self-Organizing Genome: Principles of Genome Architecture and Function. Cell 183, 28–45 (2020).

2. Rowley, M.J. & Corces, V.G. Organizational principles of 3D genome architecture. Nat Rev Genet 19, 789–800 (2018).

3. Luppino, J.M. et al. Cohesin promotes stochastic domain intermingling to ensure proper regulation of boundary-proximal genes. Nat Genet 52, 840–848 (2020).

4. Dixon, J.R. et al. Topological domains in mammalian genomes identified by analysis of chromatin interactions. Nature 485, 376–80 (2012).

5. Schuijers, J. et al. Transcriptional Dysregulation of MYC Reveals Common Enhancer-Docking Mechanism. Cell Rep 23, 349–360 (2018).

6. Liu, Y. & Dekker, J. CTCF-CTCF loops and intra-TAD interactions show differential dependence on cohesin ring integrity. Nat Cell Biol 24, 1516–1527 (2022).

7. Hansen, A.S. CTCF as a boundary factor for cohesin-mediated loop extrusion: evidence for a multi-step mechanism. Nucleus 11, 132–148 (2020).

8. Ortabozkoyun, H. et al. CRISPR and biochemical screens identify MAZ as a cofactor in CTCF-mediated insulation at Hox clusters. Nat Genet 54, 202–212 (2022).

9. Kane, L. et al. Cohesin is required for long-range enhancer action at the Shh locus. Nat Struct Mol Biol 29, 891–897 (2022).

10. Lupianez, D.G. et al. Disruptions of topological chromatin domains cause pathogenic rewiring of gene-enhancer interactions. Cell 161, 1012–1025 (2015).

11. Chakraborty, S. et al. Enhancer-promoter interactions can bypass CTCF-mediated boundaries and contribute to phenotypic robustness. Nat Genet 55, 280–290 (2023).

12. Collins, P.L. et al. DNA double-strand breaks induce H2Ax phosphorylation domains in a contact-dependent manner. Nat Commun 11, 3158 (2020).

13. Cremer, M. et al. Cohesin depleted cells rebuild functional nuclear compartments after endomitosis. Nat Commun 11, 6146 (2020).

14. Sima, J. et al. Identifying cis Elements for Spatiotemporal Control of Mammalian DNA Replication. Cell 176, 816–830 e18 (2019).

15. Klein, K.N. et al. Replication timing maintains the global epigenetic state in human cells. Science 372, 371–378 (2021).

16. Abramo, K., Valton, A.L., Venev, S.V., Ozadam, H., Fox, A.N. & Dekker, J. A chromosome folding intermediate at the condensin-to-cohesin transition during telophase. Nat Cell Biol 21, 1393–1402 (2019).

17. Emerson, D.J. et al. Cohesin-mediated loop anchors confine the locations of human replication origins. Nature 606, 812–819 (2022).

18. Finn, E.H. et al. Extensive Heterogeneity and Intrinsic Variation in Spatial Genome Organization. Cell 176, 1502–1515 e10 (2019).

19. Gabriele, M. et al. Dynamics of CTCF- and cohesin-mediated chromatin looping revealed by live-cell imaging. Science 376, 496–501 (2022).

20. Dekker, J. & Mirny, L.A. The chromosome folding problem and how cells solve it. Cell 187, 6424–6450 (2024).

21. Uhlmann, F. A unified model for cohesin function in sisterchromatid cohesion and chromatin loop formation. Mol Cell 85, 1058–1071 (2025).

22. Fujishiro, S., Sasai, M. & Maeshima, K. Chromatin domains in the cell: Phase separation and condensation. Curr Opin Struct Biol 91, 103006 (2025).

23. Alonso-Gil, D. & Losada, A. NIPBL and cohesin: new take on a classic tale. Trends Cell Biol 33, 860–871 (2023).

24. Luppino, J.M. et al. Co-depletion of NIPBL and WAPL balance cohesin activity to correct gene misexpression. PLoS Genet 18, e1010528 (2022).

25. Casa, V. et al. Redundant and specific roles of cohesin STAG subunits in chromatin looping and transcriptional control. Genome Res 30, 515–527 (2020).

26. Kojic, A. et al. Distinct roles of cohesin-SA1 and cohesin-SA2 in 3D chromosome organization. Nat Struct Mol Biol 25, 496–504 (2018).

27. Rolef Ben-Shahar, T., et al. Eco1-dependent cohesin acetylation during establishment of sister chromatid cohesion. Science 321, 563–6 (2008).

28. Minamino, M. et al. Esco1 Acetylates Cohesin via a Mechanism Different from That of Esco2. Curr Biol 25, 1694–706 (2015).

29. Wutz, G. et al. ESCO1 and CTCF enable formation of long chromatin loops by protecting cohesin(STAG1) from WAPL. Elife 9(2020).

30. Park, D.S. et al. High-throughput Oligopaint screen identifies druggable 3D genome regulators. Nature 620, 209–217 (2023).

31. Sole-Ferran, M. & Losada, A. Cohesin in 3D: development, differentiation, and disease. Genes Dev 39, 679–696 (2025).

32. MacPherson, M.J., Beatty, L.G., Zhou, W., Du, M. & Sadowski, P.D. The CTCF insulator protein is posttranslationally modified by SUMO. Mol Cell Biol 29, 714–25 (2009).

33. Farrar, D. et al. Mutational analysis of the poly(ADP-ribosyl)ation sites of the transcription factor CTCF provides an insight into the mechanism of its regulation by poly(ADP-ribosyl)ation. Mol Cell Biol 30, 1199–216 (2010).

34. Yu, W. et al. Poly(ADP-ribosyl)ation regulates CTCF-dependent chromatin insulation. Nat Genet 36, 1105–10 (2004).

35. Del Rosario, B.C. et al. Exploration of CTCF post-translation modifications uncovers Serine-224 phosphorylation by PLK1 at pericentric regions during the G2/M transition. Elife 8(2019).

36. Rao, S.S.P. et al. Cohesin Loss Eliminates All Loop Domains. Cell 171, 305–320 e24 (2017).

37. Scholz, B.A. et al. WNT signaling and AHCTF1 promote oncogenic MYC expression through super-enhancer-mediated gene gating. Nat Genet 51, 1723–1731 (2019).

38. Lancho, O. & Herranz, D. The MYC Enhancer-ome: Long-Range Transcriptional Regulation of MYC in Cancer. Trends Cancer 4, 810–822 (2018).

39. Finn, E.H. & Misteli, T. A high-throughput DNA FISH protocol to visualize genome regions in human cells. STAR Protoc 2, 100741 (2021).

40. Shachar, S., Voss, T.C., Pegoraro, G., Sciascia, N. & Misteli, T. Identification of Gene Positioning Factors Using High-Throughput Imaging Mapping. Cell 162, 911–23 (2015).

41. Almansour, F., Keikhosravi, A., Pegoraro, G. & Misteli, T. Allele-level visualization of transcription and chromatin by high-throughput imaging. Histochem Cell Biol 162, 65–77 (2024).

42. Keikhosravi, A. et al. High-throughput image processing software for the study of nuclear architecture and gene expression. Sci Rep 14, 18426 (2024).

43. Haarhuis, J.H.I. et al. The Cohesin Release Factor WAPL Restricts Chromatin Loop Extension. Cell 169, 693–707 e14 (2017).

44. Benedict, B. et al. WAPL-Dependent Repair of Damaged DNA Replication Forks Underlies Oncogene-Induced Loss of Sister Chromatid Cohesion. Dev Cell 52, 683–698 e7 (2020).

45. Kavli, B. et al. RPA2 winged-helix domain facilitates UNG-mediated removal of uracil from ssDNA; implications for repair of mutagenic uracil at the replication fork. Nucleic Acids Res 49, 3948–3966 (2021).

46. Marechal, A. & Zou, L. RPA-coated single-stranded DNA as a platform for post-translational modifications in the DNA damage response. Cell Res 25, 9–23 (2015).

47. Mishra, A.K., Dormi, S.S., Turchi, A.M., Woods, D.S. & Turchi, J.J. Chemical inhibitor targeting the replication protein A-DNA interaction increases the efficacy of Pt-based chemotherapy in lung and ovarian cancer. Biochem Pharmacol 93, 25–33 (2015).

48. VanderVere-Carozza, P.S. et al. In Vivo Targeting Replication Protein A for Cancer Therapy. Front Oncol 12, 826655 (2022).

49. Gavande, N.S., VanderVere-Carozza, P.S., Pawelczak, K.S., Vernon, T.L., Jordan, M.R. & Turchi, J.J. Structure-Guided Optimization of Replication Protein A (RPA)-DNA Interaction Inhibitors. ACS Med Chem Lett 11, 1118–1124 (2020).

50. Toledo, L.I. et al. ATR prohibits replication catastrophe by preventing global exhaustion of RPA. Cell 155, 1088–103 (2013).

51. Glanzer, J.G., Liu, S., Wang, L., Mosel, A., Peng, A. & Oakley, G.G. RPA inhibition increases replication stress and suppresses tumor growth. Cancer Res 74, 5165–72 (2014).

52. Jang, S.W. & Kim, J.M. The RPA inhibitor HAMNO sensitizes Fanconi anemia pathway-deficient cells. Cell Cycle 21, 1468–1478 (2022).

53. Feng, Y. et al. Targeting RPA promotes autophagic flux and the antitumor response to radiation in nasopharyngeal carcinoma. J Transl Med 21, 738 (2023).

54. Dueva, R. et al. Chemical Inhibition of RPA by HAMNO Alters Cell Cycle Dynamics by Impeding DNA Replication and G2-to-M Transition but Has Little Effect on the Radiation-Induced DNA Damage Response. Int J Mol Sci 24(2023).

55. Flury, V. et al. Recycling of modified H2A-H2B provides short-term memory of chromatin states. Cell 186, 1050–1065 e19 (2023).

56. Petryk, N. et al. MCM2 promotes symmetric inheritance of modified histones during DNA replication. Science 361, 1389–1392 (2018).

57. Petryk, N. et al. Replication landscape of the human genome. Nat Commun 7, 10208 (2016).

58. Marchal, C., Sima, J. & Gilbert, D.M. Control of DNA replication timing in the 3D genome. Nat Rev Mol Cell Biol 20, 721–737 (2019).

59. Miura, H., Takahashi, S., Poonperm, R., Tanigawa, A., Takebayashi, S.I. & Hiratani, I. Single-cell DNA replication profiling identifies spatiotemporal developmental dynamics of chromosome organization. Nat Genet 51, 1356–1368 (2019).

60. Pope, B.D. et al. Topologically associating domains are stable units of replication-timing regulation. Nature 515, 402–5 (2014).

61. Balzarotti, F. et al. Nanometer resolution imaging and tracking of fluorescent molecules with minimal photon fluxes. Science 355, 606–612 (2017).

62. Scheiderer, L., Marin, Z. & Ries, J. MINFLUX achieves molecular resolution with minimal photons. Nat Photonics 19, 238–247 (2025).

63. Ostersehlt, L.M. et al. DNA-PAINT MINFLUX nanoscopy. Nat Methods 19, 1072–1075 (2022).

64. Guin, K., Keikhosravi, A., Chari, R., Pegoraro, G. & Misteli, T. Orderly mitosis shapes interphase genome architecture. Elife 14(2026).

65. Fazel, M. et al. High Resolution Fluorescence Lifetime Maps from Minimal Photon Counts. ACS Photonics 9, 1015–1025 (2022).

66. Fazel, M. et al. High-precision estimation of emitter positions using Bayesian grouping of localizations. Nat Commun 13, 7152 (2022).

67. Zhao, P.A., Rivera-Mulia, J.C. & Gilbert, D.M. Replication Domains: Genome Compartmentalization into Functional Replication Units. Adv Exp Med Biol 1042, 229–257 (2017).

68. Murayama, Y., Samora, C.P., Kurokawa, Y., Iwasaki, H. & Uhlmann, F. Establishment of DNA-DNA Interactions by the Cohesin Ring. Cell 172, 465–477 e15 (2018).

69. Tan, J. & Martin, S.E. Validation of Synthetic CRISPR Reagents as a Tool for Arrayed Functional Genomic Screening. PLoS One 11, e0168968 (2016).

70. Natsume, T., Kiyomitsu, T., Saga, Y. & Kanemaki, M.T. Rapid Protein Depletion in Human Cells by Auxin-Inducible Degron Tagging with Short Homology Donors. Cell Rep 15, 210–218 (2016).

71. Pachitariu, M. & Stringer, C. Cellpose 2.0: how to train your own model. Nat Methods 19, 1634–1641 (2022).

72. Team, R.C. R: A Language and Environment for Statistical Computing. Vol. 4.3.3 (R Foundation for Statistical Computing, Vienna, Austria, 2024).

73. Boutros, M., Bras, L.P. & Huber, W. Analysis of cell-based RNAi screens. Genome Biol 7, R66 (2006).

74. Schnitzbauer, J., Strauss, M.T., Schlichthaerle, T., Schueder, F. & Jungmann, R. Super-resolution microscopy with DNA-PAINT. Nat Protoc 12, 1198–1228 (2017).

75 Kim, M., et al. Multifaceted roles of cohesin in regulating transcriptional loops. bioRxiv (2024).

